# Spinal cord synaptic plasticity by GlyRβ release from receptor fields and syndapin-dependent uptake

**DOI:** 10.1101/2021.08.18.456917

**Authors:** Jessica Tröger, Eric Seemann, Rainer Heintzmann, Michael M. Kessels, Britta Qualmann

**Author notes:** Correspondence &.

## Abstract

Glycine receptor-mediated inhibitory neurotransmission is key for spinal cord function. Recent observations suggested that by largely elusive mechanisms also glycinergic synapses display synaptic plasticity. We here identify syndapin I as critical player. Interestingly, syndapin I cooperates but in part also competes with gephyrin. Syndapin I deficiency led to fragmentation of glycine receptor fields, more disperse receptors and increased receptor mobility. Kainate treatment highlighted syndapin I’s importance even more. Our analyses unveiled that PKC-mediated S403 phosphorylation-mediated glycine receptor β decoupling from gephyrin scaffolds simultaneously promoted syndapin I association. In line, kainate-treated syndapin I KO spinal cords showed even more severe receptor field fragmentation. Furthermore, syndapin I deficiency completely disrupted kainate-induced glycine receptor internalization. Together, this unveiled important mechanisms controlling the number and organization of glycine receptor fields at inhibitory postsynapses during both steady-state and kainate-induced synaptic rearrangement - principles organizing and fine-tuning synaptic efficacy of inhibitory synapses in the spinal cord.

## Introduction

Inhibitory neurotransmission in spinal cord and brain stem is glycine receptor (GlyR)-mediated. Postsynaptic GlyRs are heteromeric pentamers containing α_1_-α_4_ and β subunits, whereas homomeric GlyRs (5 α-subunits) can also be found as extrasynaptic reservoir (Legendre, 2002, Dutertre et al, 2012). Synaptic receptor arrays are formed by the **s**caffolding protein gephyrin (Kirsch & Betz, 1995), which interacts tightly with β-subunits (GlyRβ) (Grudzinska et al, 2005, Dumoulin et al, 2010, Alvarez, 2017, Kasaragod & Schindelin, 2018.

Synaptic strength can be modulated by changes of the amount of neurotransmitter receptors in the postsynaptic membrane. Whereas mechanisms of synaptic plasticity are well understood for glutamate receptors in excitatory synapses in the hippocampus (Diering & Huganir, 2018, Choquet & Hosy, 2020), mechanisms that modulate the activity of inhibitory synapses in spinal cord and brain stem are far less understood. The activation of N-methyl-D-aspartate (NMDA) receptors was observed to decrease lateral GlyR diffusion, to increase GlyR cluster number and to correspondingly result in higher amplitudes of glycinergic miniature inhibitory postsynaptic currents (mIPSCs) (Lévi et al., 2008). In contrast, activation of protein kinase C (PKC), for example via kainate receptor stimulation, led to i) increased lateral GlyR diffusion, ii) decreased GlyR abundance at inhibitory synapses and iii) S403 phosphorylation in the cytoplasmic loop of GlyRβ reducing gephyrin binding (Specht et al., 2011). Kainate stimulation also led to GlyR endocytosis (Sun et al., 2014). Also this modulation of receptor surface availability was reported to be PKC-dependent. It resulted in dramatic reductions of GlyR puncta in cultured spinal cord neurons (Sun et al., 2014). Yet, apart from the apparently required decoupling from gephyrin, the organizational changes of inhibitory receptor arrays and components that are critical for GlyR dynamics largely remained elusive.

The large intracellular loop of GlyRβ binds to the membrane-binding F-BAR protein syndapin I (also called PACSIN1) (Qualmann et al, 1999, Qualmann et al, 2011, Kessels & Qualmann, 2015) at a site distinct from, but adjacent to, the gephyrin binding site and syndapin I deficiency was reported to cause some reduction of GlyR puncta (del Pino et al, 2014). A GlyRα mutation, which coincided with reduced syndapin I interaction in in vitro-reconstitution screenings with GlyR peptides, was accompanied by startle disease and showed disturbed glycinergic neurotransmission (Langlhofer et al, 2020).

Here we demonstrate by (ultra-)high resolution analyses of spinal cords that syndapin I knock-out (KO) does not lead to a reduction of GlyRβ at the plasma membrane but to fragmentations of GlyR fields. Syndapin I deficiency also increased GlyRβ mobility. Under kainate stimulation, the pivotal role of syndapin I in the organization of GlyR fields and in the modulation of GlyR availability became even more visible. Regulatory mechanisms, which decouple GlyRβ subunits from gephyrin, promoted GlyR/syndapin I interactions. Furthermore, syndapin I was identified as crucial for GlyR internalization during kainate-induced reorganization.

Our work highlights a critical element in glycine receptor field organization and in the thus far poorly understood kainate-induced GlyRβ dynamics of inhibitory synapses.

## Results

### Binding of syndapin I and gephyrin to the cytoplasmic loop of GlyRβ is not mutually exclusive but shows partial competition

Using adjacent sites in the large cytoplasmic loop, GlyRβ interacts with both gephyrin and the membrane-binding protein syndapin I (del Pino et al, 2014). This raised the question whether these two proteins act independently in regulating GlyRs by competing with each other for binding or whether syndapin I and gephyrin may show some cooperative functions. Since syndapin I did not associate with gephyrin (**Figure 1A,B**), putative complex formations of syndapin I with gephyrin would be indirect and GlyRβ-mediated.

**Figure 1.**
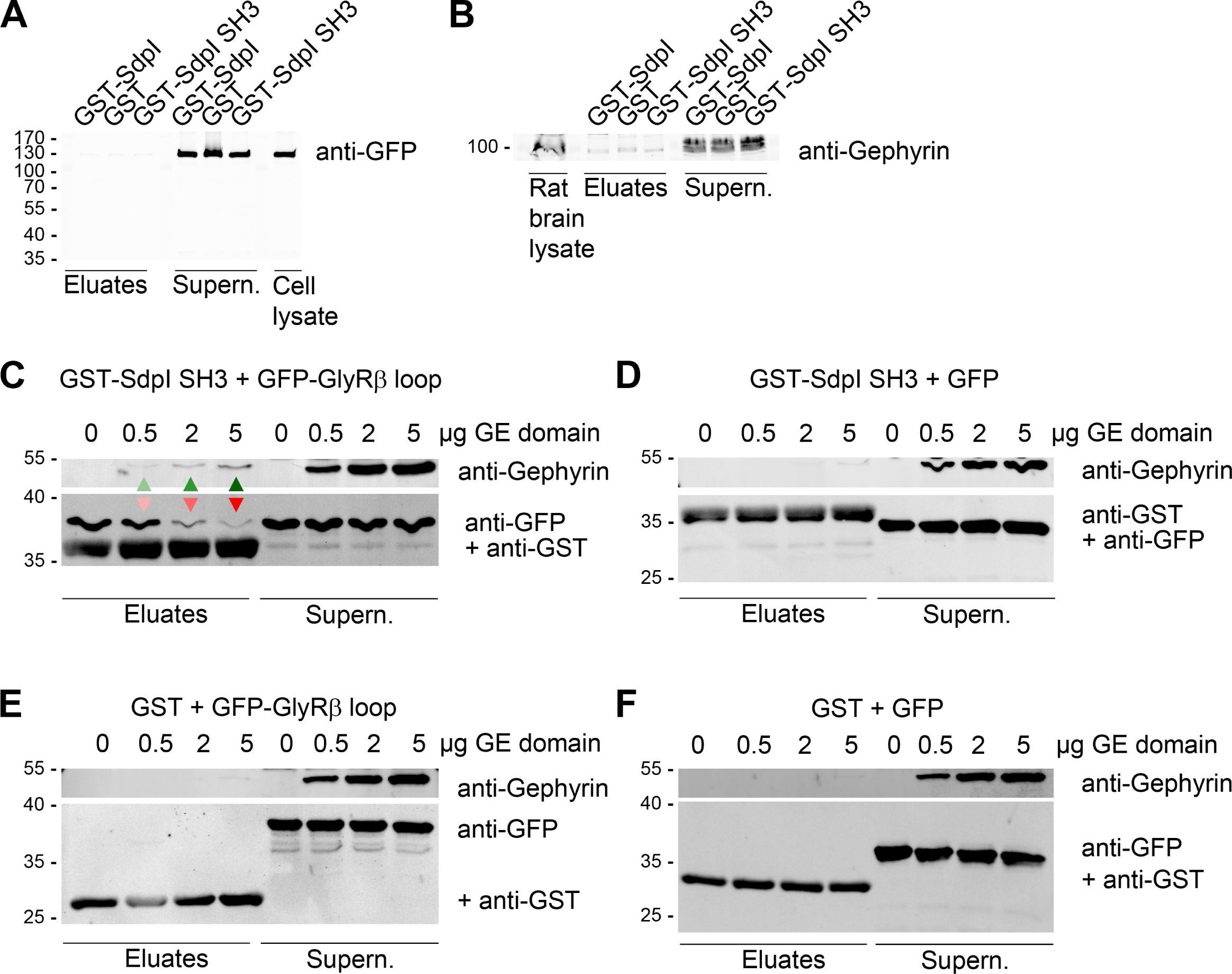
Binding of syndapin I and gephyrin to the cytoplasmic loop of GlyRβ is not mutually exclusive but show partial competition. (**A,B**) Immunoblotting analyses of attempts to coprecipitate GFP-gephyrin expressed in HEK293 cells (**A**) and endogenous gephyrin from rat brain lysates (**B**) with immobilized GST-syndapin I (GST-SdpI) and GST-syndapin I SH3 domain (GST-SdpI SH3) and GST (negative control) showing that syndapin I does not bind to gephyrin. Gephyrin remained in the supernatants in both experiments. (**C-F**) Demonstration of complexes composed of GST-syndapin I SH3 domain (GST-SdpI SH3), GFP-GlyRβ cytoplasmic loop (GFP-GlyRβ loop) and gephyrin E domain (GE domain) by specific coprecipitation of gephyrin E domain by immobilized GST-SdpI SH3 (**C,D**) but not by GST (**E,F**) in the presence of GFP- GlyRβ loop (**C**) but not in the absence of GlyRβ E-loop (GFP control; **D**). Green arrowheads mark rising amounts of GE domain bound. Red arrowheads mark the decrease of GST-SdpI SH3-associated GFP- GlyRβ cytoplasmic loop upon rising amounts of GE domain added. Besides this competition, however, note that especially at the higher concentrations of GE domain tested, complexes composed of all three binding partners (SdpI SH3, GlyRβ loop and GE domain) were successfully formed (arrowheads) (**C**).

Reconstitutions of complex formation with purified GST-syndapin I SH3 domain, GFP-tagged GlyRβ’s cytoplasmic loop expressed in HEK293 cells and purified, GlyRβ-binding gephyrin E domain showed that increasing amounts of gephyrin E domain diminished the amount of syndapin I SH3 domain-associated GlyRβ (**Figure 1C**). Thus, there was considerable steric competition between the two different GlyRβ binding partners.

Yet, GFP-tagged GlyRβ cytoplasmic loop coprecipitated by immobilized syndapin I SH3 domain obviously also allowed for simultaneous association of some gephyrin E domain, i.e. complexes of all three proteins were formed, too (**Figure 1C-F**; arrowheads).

Taken together, the GlyRβ binding of syndapin I and gephyrin showed some - probably steric – competition but clearly was not mutually exclusive.

### Increased microscopic resolution revealed that syndapin I KO does not lead to reduced but increased density of smaller GlyRβ-containing receptor clusters

To shed light on a putative common, cooperative role of syndapin I and gephyrin in GlyR-mediated neurotransmission, we first focused on receptor scaffolding functions represented by gephyrin.

Quantitative analyses of spinal cord homogenates showed that syndapin I KO did not alter the expression of GlyRβ when compared to WT (**Figure 2A,B**). This result was hard to reconcile with previous immunofluorescence analyses suggesting a reduced GlyR cluster density (about -20%) upon syndapin I KO using a pan anti-GlyR antibody (del Pino et al, 2014). Intriguingly, structured illumination microscopy (SIM) unveiled that syndapin I KO did in fact not reduce the number of GlyRβ clusters along dendrites in comparison to wild-type (WT) neurons but instead increased them (**Figure 2C,D; Figure EV1A-D**). The GlyRβ cluster density rose by about 27% (**Figure 2E**). Additionally, GlyRβ clusters of syndapin I KO spinal cord neurons were smaller than those of WT neurons (**Figure 2F**).

**Figure 2.**
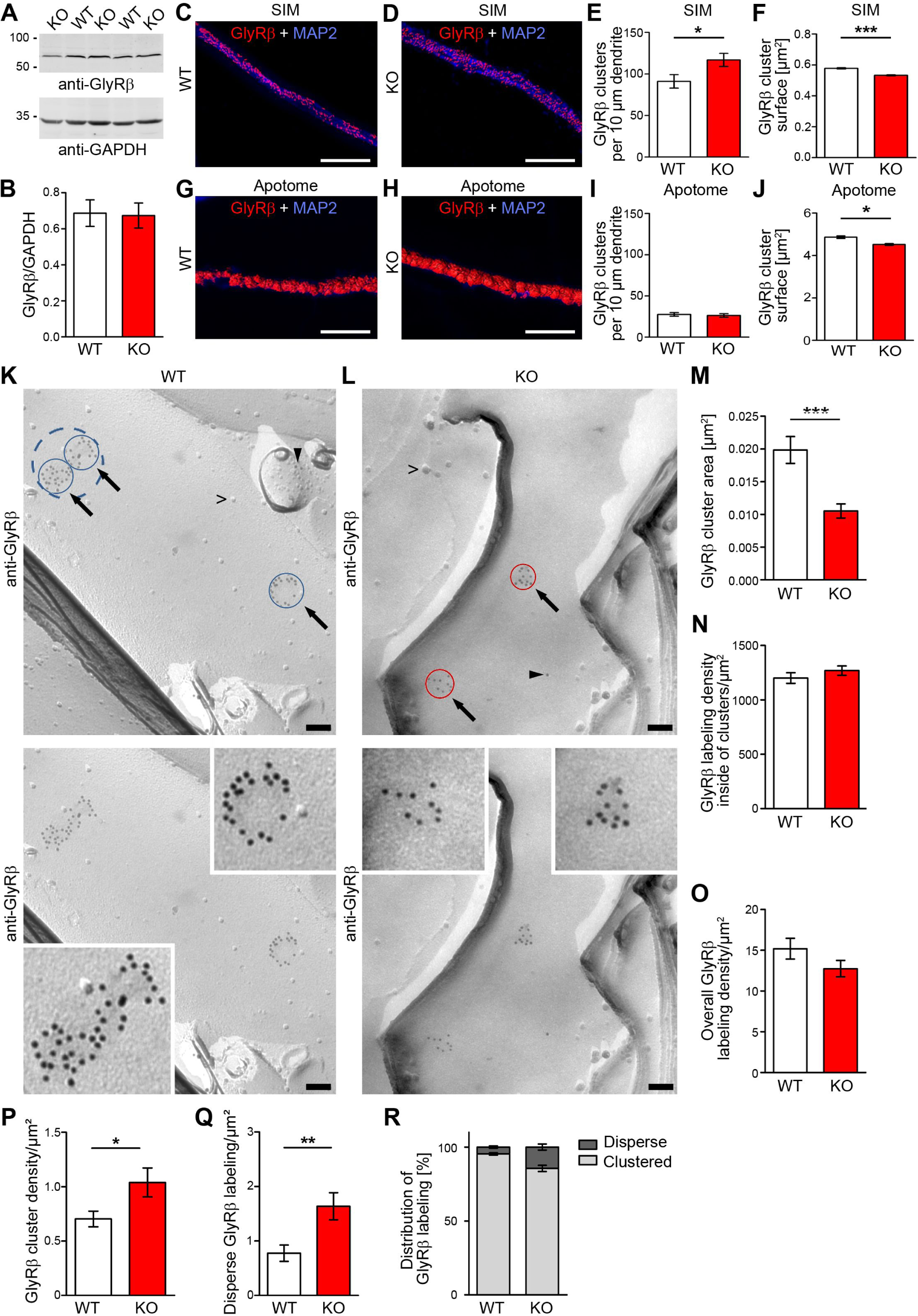
High-resolution analyses unveil an increased abundance and a decreased size of GlyRβ clusters upon syndapin I KO. (**A,B**) Quantitative immunoblotting analyses of GlyRβ expression levels in WT and syndapin I KO spinal cord homogenates (normalized to anti-GAPDH levels; homogenates of n=10 mice/genotype). (**C-J**) Analyses of GlyRβ clusters along dendrites of spinal cord neurons (DIV21/22) from WT (**C,G**) and syndapin I KO mice (**D,H**). Shown are 3D reconstructions of GlyRβ clusters (red) overlaid on anti- MAP2 immunolabeled dendritic areas (blue) using images recorded by structured illumination microscopy (SIM) (**C,D**) and conventional light microscopy (Apotome-assisted) (**G,H)**. Single channels for comparisons of microscopy image versus 3D-reconstructions see **Figure EV1A-H**. Quantitative analyses of cluster densities (**E,I**) and sizes (**F,J**). Bars, 3 µm. **(K,L)** TEM images of freeze-fractured spinal cord plasma membranes from WT (**K**) and syndapin I KO mice (**L**) labeled with immunogold conjugates directed against the intracellular loop of the GlyRβ (pointed out by arrows). Note that the GlyRβ clusters detected are larger in WT (**K**; clusters are encircled in blue; an alternative consideration (large cluster instead of two adjacent ones) is encircled by a dashed line) than in syndapin I KO specimen (**L**; clusters encircled in red). Arrowheads mark disperse GlyRβ immunogold labelings (1 and 2 gold particles). Lower panels show the above TEM images without any putatively covering labeling. Insets show the GlyRβ clusters at enhanced magnification and increased contrast. Bars, 100 nm. (**M-R**) Quantitative analyses of GlyRβ immunolabeling at the plasma membrane in WT and syndapin I KO samples of spinal cords including determinations of the GlyRβ cluster sizes (**M**), the labeling density within receptor fields (**N**), the density of membrane-bound anti-GlyRβ immunolabeling in total (**O**), the density of GlyRβ clusters (**P**), the density of disperse anti-GlyRβ immunolabeling (**Q**) and the distribution of anti-GlyRβ immunogold labeling between clustered and disperse (**R**). Data, mean±SEM. (**E,I**) n=20 each (SIM) and n=34/36 (WT/KO) (Apotome) dendrite segments and cells per genotype of 3 independent neuron preparations. (**F,J**) n=10184/12855 (WT/KO) (**F**) and 7144/7234 (WT/KO) (**J**) GlyRβ clusters. (**M-R**) n=50/49 (WT/KO) images for density determinations and n=96/139 (WT/KO) individual GlyRβ clusters from two independent spinal cord preparations. Unpaired student’s t-test (**B,E,I**; n.s); Mann Whitney (**F,J,M-Q**). *, p<0.05; **, p<0.01; ***, p<0.001.

To address whether the gross discrepancy to literature data represented a lab-, a reagent- or experimenter-related inconsistency and/or a difference arising from the different microscopy techniques used, which may also have some major impact on the analysis of subcellular structures in the nervous system (Tröger et al, 2020), we analyzed the same samples as before by a more classical technique not providing such a high resolution (Apotome 3D imaging) (**Figure 2F-H; Figure EV1E-H**). Strikingly, conventional immunofluorescence microscopy images of the same samples failed to resolve the increased density of GlyRβ clusters but led to results different from our SIM analyses. First, the absolute number of detected GlyRβ clusters was much lower than with SIM. Less than a third of all SIM-detected clusters were detected (**Figure 2I**). Second, also the increase in GlyRβ cluster density in syndapin I KO neurons was not detected by conventional immunofluorescence microscopy (**Figure 2I**).

Thus, in line with the small dimensions of inhibitory postsynapses in spinal cord neurons (Specht et al, 2013), our evaluations demonstrated that conventional light microscopy was hampered by resolution limits. Syndapin I KO does not lead to GlyRβ clusters losses (del Pino et al, 2014) but results in a significant increase of GlyRβ cluster density (**Figure 2C-E**).

We next asked whether the SIM-based observation indicating that the increase in GlyRβ cluster density along dendrites comes at the expense of a reduced amount of GlyRβ per cluster (**Figure 2F**) would consistently be seen by examinations of the same samples by conventional light microscopy. In line with reported increases of microscopic resolution by SIM (Gustafsson, 2005), the surfaces of GlyRβ clusters resolvable by the two methods differed by about one order of magnitude (e.g. WT: 0.6 µm^2^ (SIM) versus 4.8 µm^2^ (Apotome). Yet, both methods consistently revealed that GlyRβ clusters surfaces in KO neurons were significantly smaller than those in WT neurons (**Figure 2F,J**).

Taken together these results suggested a critical role of syndapin I in organizing GlyRβ arrays, i.e. a function that seemed somewhat related to that of gephyrin.

### Quantitative ultra-high resolution analysis show that syndapin I KO spinal cords have fragmented GlyR fields and a higher abundance of dispersely localized GlyRβs

Even the increased resolution of SIM may still fail to detect even smaller GlyR clusters or single, dispersed GlyRs, which could putatively also occur upon syndapin I KO. Electron microscopy allows for resolutions of better than 1 nm, i.e. can in principle resolve single proteins or even domains thereof. Receptor field examinations free from rotational artifacts would require undisturbed perpendicular views onto the plasma membrane. Although to our knowledge never attempted thus far, let alone directly in spinal cord tissue, this could in principle be accomplished via plasma membrane freeze-fracturing and subsequent anti-GlyRβ immunogold labeling. We finally were indeed able to i) freeze-fracture murine spinal cord tissue and to ii) furthermore label the resulting P-faces (cytosolic faces) of the freeze-fracture replica with anti-GlyRβ antibodies directed against cytosolic GlyRβ epitopes (**Figure 2K,L**; **Figure EV1I,J**).

In WT spinal cords, GlyR fields were visualized by strikingly high densities of anti-GlyRβ immunogold labels, if small enough colloidal gold was used. Some receptor fields showed more irregular shapes but many were relatively round and had diameters of ∼160 nm (**Figure 2K**). Double immunogold labeling attempts to also visualize syndapin I failed under the conditions suitable for GlyRβ labeling. Both ice surfaces but also the E-faces of fractured membranes were devoid of anti-GlyRβ labeling (**Figure EV1I,J**). This demonstrated the specificity of the anti-GlyRβ immunolabeling.

Syndapin I KO spinal cords also displayed GlyRβ fields of relatively circular shapes. Strikingly, our ultra-structural imaging resolved that syndapin I KO GlyRβ clusters were only about half the size of those observed in WT tissue (**Figure 2K-M**). As the anti-GlyRβ immunogold labeling density within the cluster areas (**Figure 2N**) and also the overall anti-GlyRβ immunogold labeling density at the plasma membrane showed no change upon syndapin I KO (**Figure 2O**), no plasma membrane-bound GlyRβs was lacking. Instead, the GlyRβs distribution differed. Syndapin I KO spinal cords showed a more than 50% higher GlyRβ cluster density when compared to WT (**Figure 2P**).

Additionally, non-clustered, i.e. disperse GlyRβ immunolabelings were much more abundant. The density of disperse GlyRβ immunolabeling in syndapin I KO spinal cords was more than twice as high as in WT spinal cords (**Figure 2Q**). Also distribution analyses of all labels observed unveiled that syndapin I KO spinal cords displayed a strong increase in the percentage of disperse GlyRβs (**Figure 2R**).

Syndapin I thus clearly plays a scaffolding role holding together GlyRβ fields. This finding was unexpected, as gephyrin is considered as the scaffold ensuring efficient GlyRβ clustering. Instead, it seemed that syndapin I and gephyrin cooperate in receptor scaffolding functions in a non-redundant manner, i.e. that also syndapin I plays a critical role in holding together GlyRβ fields.

### Particularly small GlyRβ clusters in syndapin I KO spinal cord neurons lack the postsynaptic scaffold gephyrin

We next asked whether the fragmented and/or shrunken GlyRβ clusters in syndapin I KO mice still contain the postsynaptic scaffold gephyrin. We hypothesized that this should be the case. Given the partial steric hindrances between gephyrin and syndapin I binding to GlyRβ (**Figure 1C**), it even seemed likely that gephyrin may take over some of the docking sites of its GlyRβ binding neighbor syndapin I when syndapin I is knocked out. To our surprise, none of this was the case. Quantitative analyses of gephyrin clusters by SIM (**Figure 3A,B**) showed that the gephyrin cluster density (**Figure 3C,D**) did not fully mirror the significant increase in GlyRβ cluster abundance (compare **Figure 2**). While also gephyrin clusters were significantly smaller upon syndapin I KO (**Figure 3D**), their density only showed moderately increased values that failed to reach statistical significance despite high n numbers of dendritic segments analyzed (**Figure 3C**). A simple explanation for the apparently lacking gephyrin was that gephyrin expression levels may be reduced in syndapin I KO mice. Quantitative immunoblotting analyses, however, showed that gephyrin levels were similar in WT and syndapin I KO spinal cords (**Figure 3E,F**).

**Figure 3.**
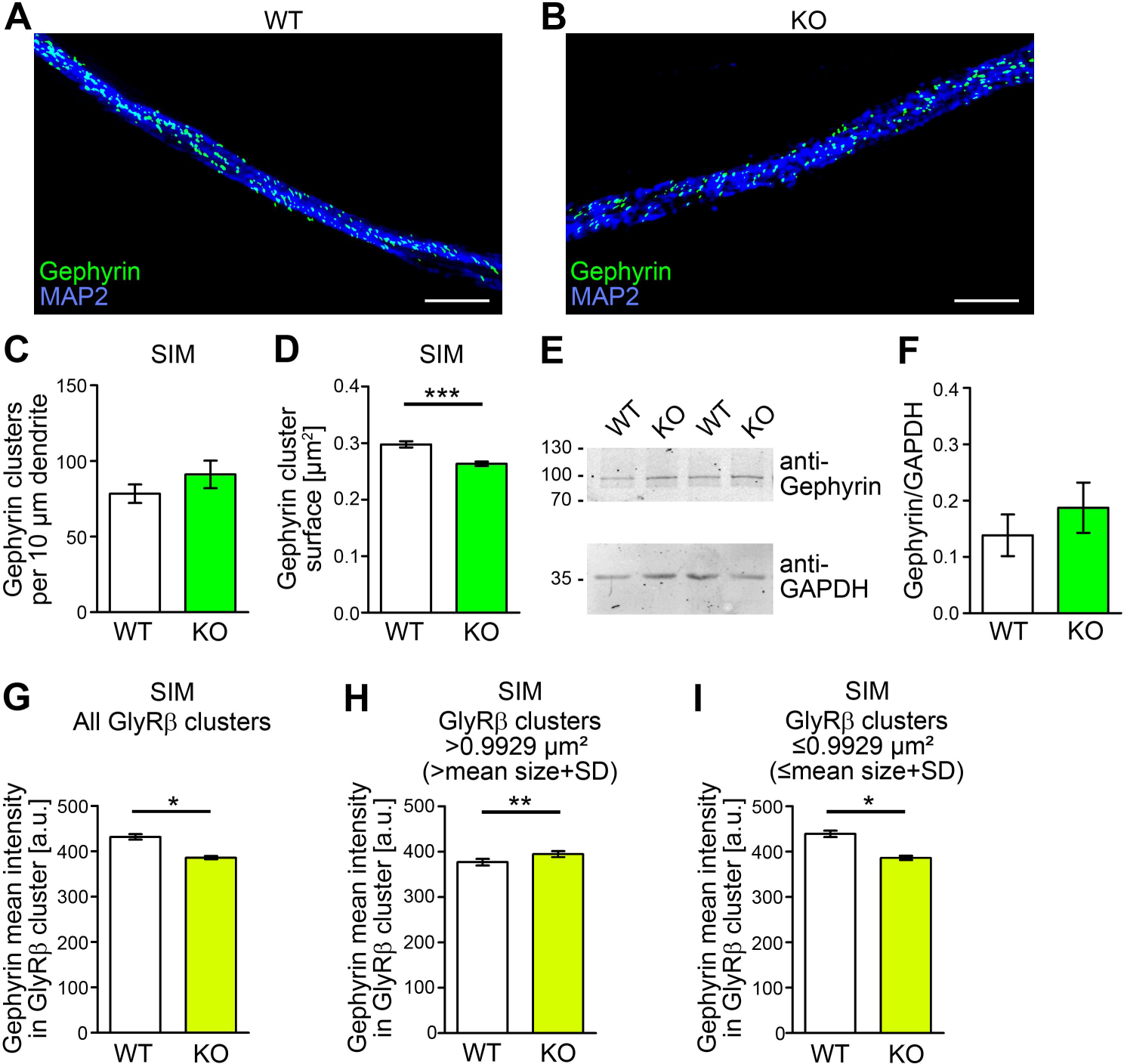
Syndapin I KO specifically reduces gephyrin clusters sizes in small GlyRβ clusters. (**A-D**) 3D reconstructions of anti-gephyrin clusters in anti-MAP2-colabeled dendrites of neurons (DIV21/22) isolated at E14 from WT (**A**) and syndapin I KO spinal cords (**B**) using SIM and quantitative analyses of the gephyrin cluster density along dendrites (**C**) and of the sizes of the gephyrin clusters (**D**). (**E,F**) Quantitative immunoblotting analyses of gephyrin expression levels in WT and syndapin I KO spinal cord homogenates (normalized to anti-GAPDH levels). (**G-I**) Quantitative analyses of mean anti-gephyrin intensities in GlyRβ clusters in dendrites of DIV21/22 spinal cord neurons isolated from WT and syndapin I KO spinal cords and imaged by SIM. The mean anti-gephyrin intensities are shown irrespective of the corresponding GlyRβ cluster size (**G**) as well as separately in i) larger (**H**) and in ii) GlyRβ clusters equal to and smaller (**I**) than the average WT GlyRβ cluster size (expressed as cluster surface of 3D-reconstructed GlyRβ clusters (0.5777 µm^2^ + standard deviation (0.4152 µm^2^), i.e. ≤0.9929 µm^2^). Data, mean±SEM. (**c**) n=21 (WT) and 23 (KO) cells of 2 independent preparations. (**D**) 9372/11982 (WT/KO) gephyrin clusters. (**E,F**) N=10 mice/genotpye. (**G-I**) n=10184/12855 (WT/KO) (**G**), 1397/1389 (WT/KO) (**H**) and 8787/11466 (WT/KO) (**I**) gephyrin clusters from 20 cells each from 3 independent preparations from 3+8+8 WT pooled embryos and 4+3+11 pooled KO embryos, respectively. Unpaired student’s t-test (**C**; n.s) and Mann Whitney (**D,F-I**), respectively. *, p<0.05; **, p<0.01; ***, p<0.001.

An alternative explanation was that a subpopulation of GlyRβ clusters in syndapin I KO neurons is devoid of detectable gephyrin and that therefore the cluster density did not seem to increase in accordance with receptor field fragmentation. Quantitative analyses of anti-gephyrin immunolabeling densities inside of GlyRβ clusters indeed demonstrated a reduction of the average anti-gephyrin immunolabeling intensity in GlyRβ clusters in syndapin I KO spinal cord neurons (**Figure 3G**). Interestingly, whereas large GlyRβ clusters in syndapin I KO neurons even showed an increase in mean anti-gephyrin labeling intensity when compared to WT (+4.6%; *P*<0.01), which could reflect an additional incorporation of gephyrins into receptor scaffolds in the absence of syndapin I, the overall gephyrin intensity reduction in GlyRβ clusters of syndapin I KO neurons was in particular occurring in GlyRβ clusters that were smaller or equal to the mean surface of GlyR clusters (including standard deviation, i.e. ≤0.9929 µm^2^) (**-**12%; *P*<0.05) (**Figure 3H,I**).

These results for the small GlyRβ clusters in syndapin I KO neurons were stunning. The observed partial competition of gephyrin and syndapin I for binding to the cytoplasmic loop of the GlyR receptor should rather allow for an improved gephyrin binding upon syndapin I KO. Yet, the opposite was observed. A subpopulation of GlyRβ clusters in syndapin I KO spinal cords has lost some gephyrin and this subpopulation corresponds to the small clusters that occur with higher abundance upon syndapin I KO.

### Syndapin I deficiency leads to an increased mobility of GlyRβ clusters

To shed some more light on syndapin I’s crucial role in GlyRβ scaffolding, we next asked whether syndapin I might contribute to synaptic GlyRβ immobilization. Upon syndapin I knock-down in rat spinal cord neurons, both the spot mobility and the displacement length of GFP-GlyRβ were significantly increased in comparison to control (**Figure 4A-D**). The mean velocity of tracked GFP-GlyRβ spots increased by 15% in syndapin I-deficient neurons (**Figure 4C**). Additionally, the mean displacement length of tracked clusters increased by about 50% in syndapin I-deficient neurons (**Figure 4D**). Both defects observed in syndapin I-deficient neurons were highly statistically significant compared to WT and thus clearly represented syndapin I loss-of-function phenotypes in receptor anchoring (**Figure 4C,D**).

**Figure 4.**
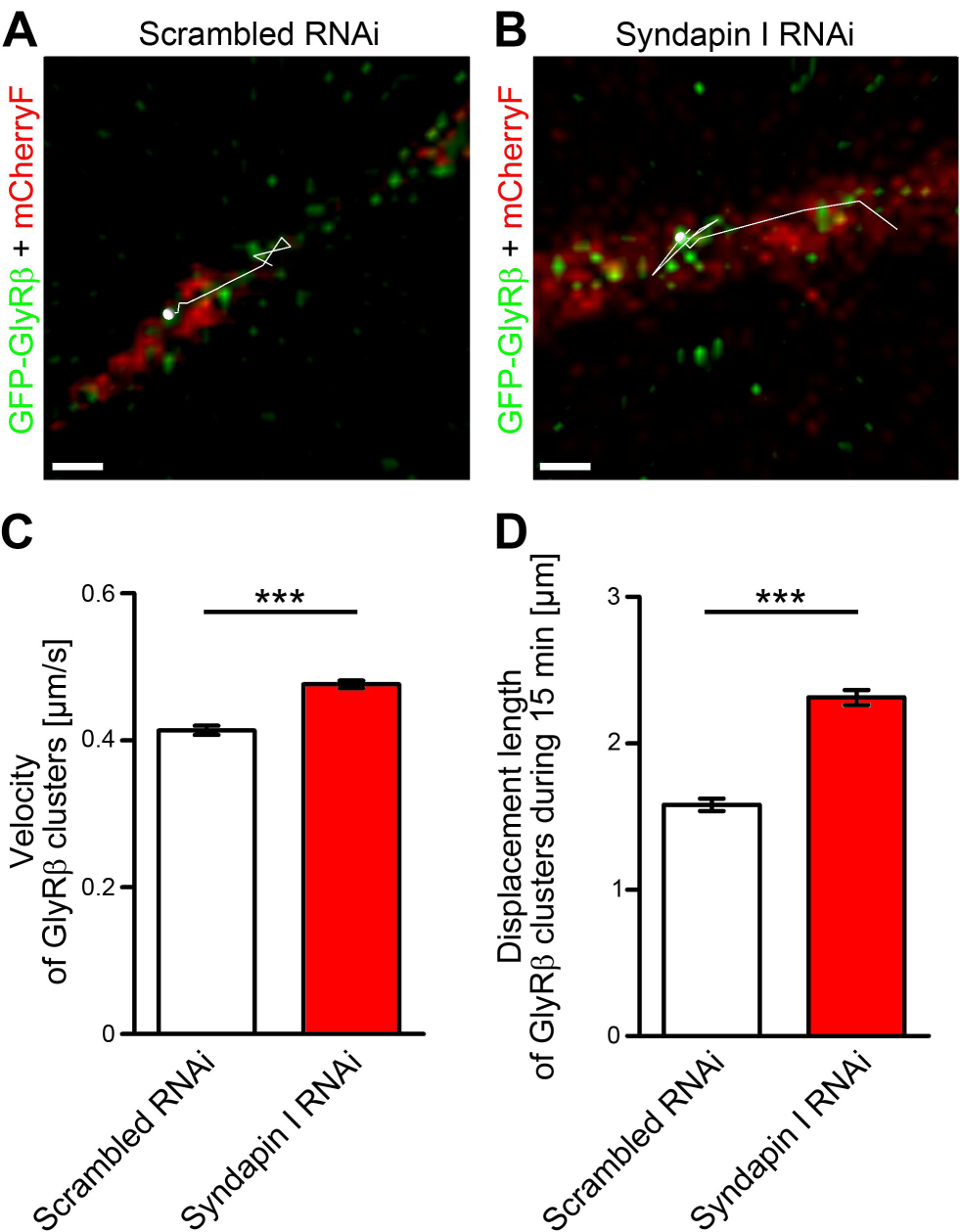
Syndapin I KO leads to an increased mobility of GFP-GlyRβ clusters. (**A,B**) Examples of trajectories from reconstructed GFP-GlyRβ clusters projected onto merged images of rat spinal cord neurons double-transfected with GFP-GlyRβ (green) and scrambled RNAi/mCherryF (red) (**A**) and with GFP-GlyRβ and syndapin I RNAi/mCherryF (**B**), respectively. Spheres show the respective start point of the GFP-GlyRβ tracking. Bar, 1 µm. (**C,D)** Quantitative analyses of GFP-GlyRβ cluster speeds (**C**) and displacement lengths (**D**) in spinal cord neurons cotransfected with either syndapin I RNAi and control (scrambled RNAi) plasmids and imaged by spinning disc microcopy. Data, mean±SEM. n=1634 tracks from 10 neurons (scrambled RNAi) and 2251 tracks from 11 neurons (syndapin I RNAi). Mann Whitney. ***, p<0.001.

### Syndapin I KO completely disrupts the kainate-induced internalization of GlyR

Biochemically, we did not only observe cooperative functions of gephyrin and syndapin I but also some competition for GlyRβ (**Figure 1**). While the defects in receptor organization observed upon syndapin I KO (**Figures 2-4**) pointed to some scaffolding role of syndapin I, which was somewhat cooperative but not redundant with gephyrin, there furthermore should be functions of syndapin I, which are promoted by the absence of gephyrin association, and thereby rather reflect the competitive behavior of gephyrin and syndapin I. In recent years, it has been shown that certain conditions lead to detectable GlyR endocytosis. PKC-induced GlyR endocytosis has been shown upon GlyR overexpression in HEK293 cells (Huang et al, 2007, Breitinger et al, 2018). Importantly, finally also the endogenous receptors in cultures of dissociated spinal cord neurons were shown to be internalized when PKC was activated via stimulating the neurons with kainate (Sun et al, 2014). The mechanisms of GlyR endocytosis and how the to-be- internalized receptors could be decoupled from the extended postsynaptic gephyrin-based scaffold holding together the large receptor fields largely remained elusive. This prompted us to hypothesize that syndapin I could represent a critical player in this process and may be involved in a molecular switching mechanism between receptor anchoring and scaffolding on one side and receptor endocytosis on the other side. Using antibody internalization assays based on the same pan-anti-GlyR antibody as published (Sun et al, 2014), we were able to reproduce the finding that almost 30% of the GlyRs were internalized upon incubating WT neurons with kainate (**Figure 5A-C**).

**Figure 5.**
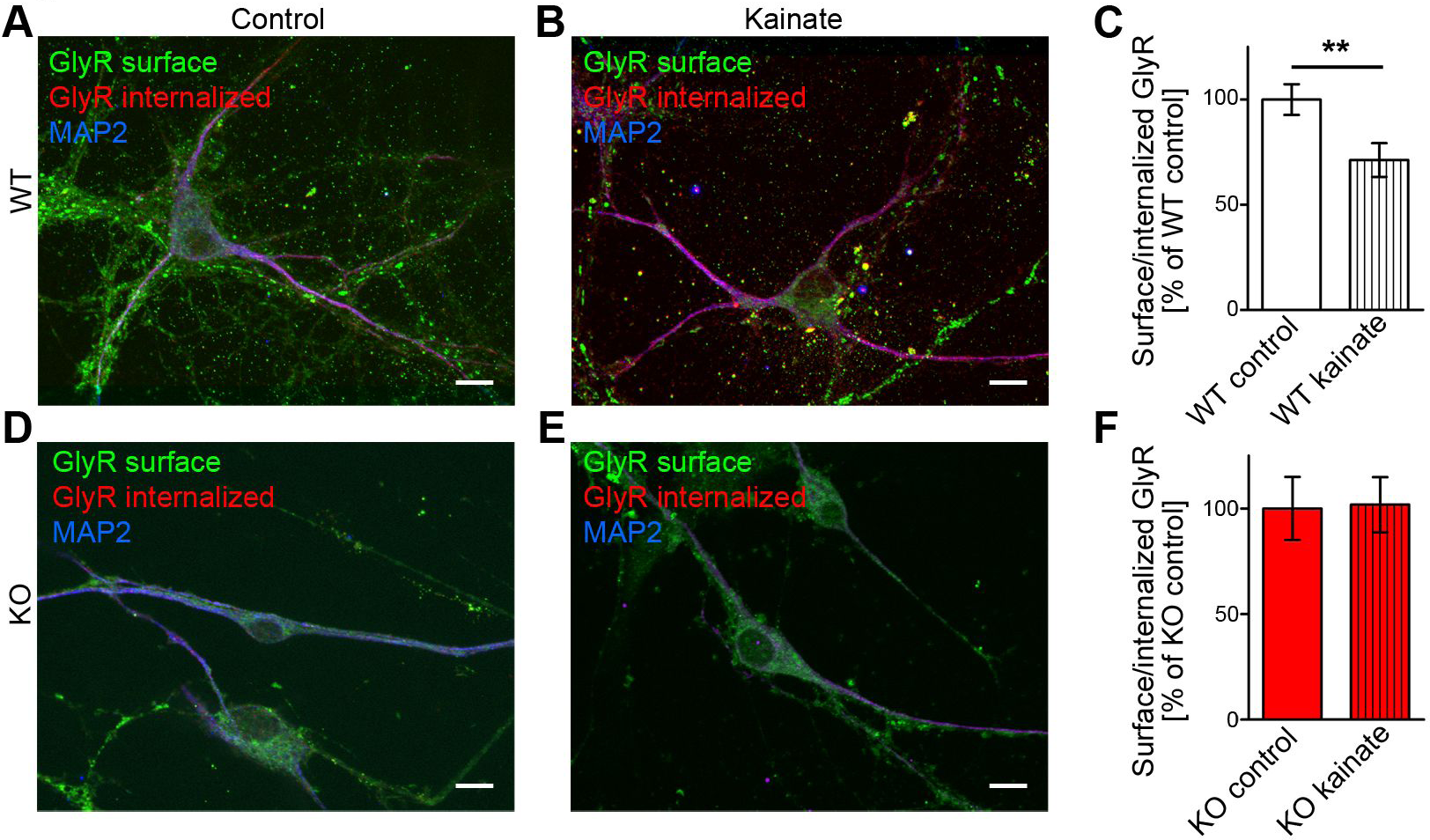
Syndapin I is crucial for GlyR endocytosis. (**A,B,D,E**) MIPs of merged immunofluorescence images (Apotome; 0.3 µm intervals) of anti-GlyR antibody internalization assays showing a internalized GlyRs (red) in WT (**A,B**) and a full block of GlyR internalization in syndapin I KO spinal cord neurons (**D,E**) when neurons were stimulated with 200 µM kainate for 1 min followed by an incubation for 10 min to allow for GlyR internalization (red). GlyR at the cell surface is shown in green, MAP2 in blue. Bars, 10 µm. Quantitative analyses (**C,F)** represent means±SEM of n=66/46 (control/kainate) (**C**) and 50/50 (control/kainate) (**F**) neurites from 3 independent experiments each. Mann Whitney. **, p<0.001.

Intriguingly, syndapin I KO completely abolished this kainate-induced GlyR endocytosis in cultures of spinal cord neurons (**Figure 5D-F**). The rates of cell surface-localized to internalized endogenous GlyR in syndapin I KO neurons did not show any response to kainate stimulation at all (**Figure 5F**).

These experiments thus identify syndapin I as critical component for kainate signaling-induced endocytosis of GlyRs in spinal cord neurons.

### Ultra-high resolutions analyses of kainate-induced receptor dynamics in spinal cords

It remained to be addressed whether the kainate-induced internalization of GlyR in WT neuronal cultures (Sun et al., 2014; this study, **Figure 5**) also occurs in intact spinal cords. Also, it is a pressing question, which type of changes in receptor field organization bring about this form of receptor and synapse modulation.

Freeze-fracture replica of kainate-stimulated spinal cords of WT mice showed a substantial decline of general GlyRβ labeling density. In contrast, in syndapin I KO spinal cords, the density of plasma membrane-localized GlyRβ did not change at all upon kainate incubation (**Figure 6A-F**). These results showed at ultra-high resolution that kainate-induced internalization of GlyRβ also occurs at the spinal cord tissue level and that syndapin I KO spinal cords are indeed completely impaired for GlyRβ internalization.

**Figure 6.**
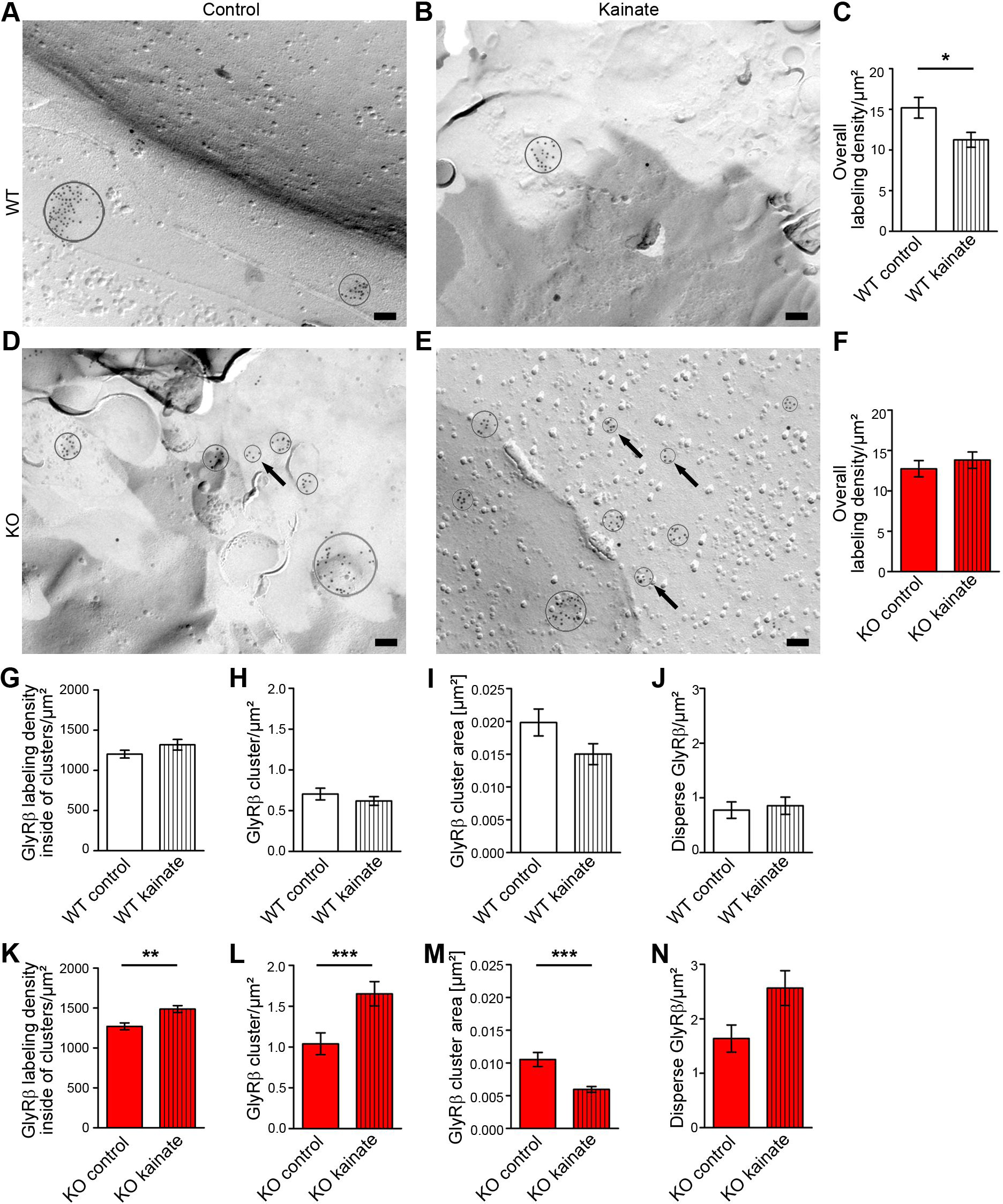
Kainate-induced synaptic plasticity in WT and syndapin I KO spinal cords leads to reorganizations of GlyRβ fields and highlights the critical role of syndapin I in GlyRβ scaffolding and internalization. **(A-F)** TEM images of anti-GlyRβ immunogold-labeled, freeze-fractured spinal cord plasma membranes from WT (**A,B**) and syndapin I KO mice (**D,E**), that were either left untreated (**A,D**) or were treated with 1 mM kainate for 60 min (**B,E**) and quantitative analyses of the overall GlyRβ immunolabeling densities thereof (**C,F**). Clusters are encircled. Note that in average clusters are smaller when syndapin I KO spinal cords were incubated with kainate (particularly small clusters are pointed out with arrows in both syndapin I KO control (**D**) and in kainate-treated syndapin I KO spinal cord (**E**). Bars, 100 nm. (**G-N**) Quantitative analyses of GlyRβ immunolabeling at the plasma membrane in WT (**G-J**) and syndapin I KO (**K-N**) samples addressing changes in receptor field organization and GlyRβ distribution upon kainate treatment. Shown are determinations of the GlyRβ labeling density inside of clusters (**G,K**), GlyRβ cluster densities (**H,L**), GlyRβ cluster sizes (**I,M**) and the densities of disperse anti-GlyRβ immunolabelings (**J,N**). Data, mean±SEM. n=50 (WT, unstimulated, see also Figure 2), n=49 (KO, unstimulated, see also Figure 2), n=45 (WT, stimulated) and n=53 (KO, stimulated) images as well as n=96 (WT, unstimulated), n=139 (KO, unstimulated), n=76 (WT, stimulated) and n=239 (KO, stimulated) individual GlyRβ clusters, respectively, from two independent spinal cord preparations. Mann Whitney (**C,F,G-N**). *, p<0.05; **, p<0.01***, p<0.001.

What may be the receptor reorganizations allowing for the kainate-induced disappearance of a significant proportion of GlyRβs from the plasma membrane in WT spinal cords? It was conceivable that GlyRβ fields would become more loosely connected due to the kainate- and PKC-mediated decrease of gephyrin binding. Yet, we did not observe that receptor packaging inside of receptor fields showed any signs of decrease upon kainate, rather the average packaging density increased (**Figure 6G**; n.s.). It thus appeared likely that receptor fields may simply break into smaller fragments upon stimulating spinal cords with kainate. Yet, our quantitative analyses showed that the GlyRβ cluster density along dendrites did not increase but again rather showed the opposite trend (**Figure 6H**; n.s.).

How do then more than 25% of all GlyRβs disappear from the plasma membrane upon kainate incubation (**Figure 6C**)? We thus next analyzed whether individual receptors may leave the receptor fields. As the overall packaging of receptors within the fields was unchanged (**Figure 6G**), such a mechanism should manifest in smaller receptor field areas and in an increase of disperse GlyRβ immunolabeling at the plasma membrane. We determined a decline of the receptor field sizes by about 25% on average. However, this decline failed to reach statistical significance despite our efforts analyzing 96 (control) and 76 (kainate) anti-GlyRβ clusters (**Figure 6I**; n.s.). A statistically significant increase of GlyRβs localized in a disperse manner in the plasma membrane was not observed either (**Figure 6J**; n.s.).

We thus concluded that the reorganizations of the receptor fields upon kainate stimulation in WT spinal cords involve a combination of receptor field breakage and shrinkage and that receptors or small clusters of receptors removed from the receptor fields largely do not remain at the plasma membrane of WT spinal cords.

### Kainate-treated spinal cords of syndapin I KO mice show impaired GlyRβ uptake and a massive disruption of GlyRβ fields

Syndapin I deficiency led to an impairment of kainate-induced GlyR internalization in dissociated neurons (**Figure 5**). Similarly, a complete disruption of GlyRβ internalization was observed in spinal cords of syndapin I KO mice (**Figure 6D-F**). The overall density of GlyRβ at the plasma membrane of syndapin I KO spinal cords did not decline at all (**Figure 6F**; 13±1/µm^2^ (control) vs. 14±1/µm^2^ (kainate); n.s.).

The lack of any change in GlyRβ density at the plasma membrane may imply that there is no kainate-induced plasticity in syndapin I KO neurons. Strikingly, however, ultra-high-resolution analyses of GlyRβ at the plasma membrane revealed that this was a misassumption. The anti-GlyRβ labeling density inside of GlyRβ clusters increased (**Figure 6K**). Also the density of GlyRβ clusters in syndapin I KO spinal cords, which was already elevated when compared to WT (**Figure 2P**), increased even further when kainate was added (**Figure 6L**). Correspondingly, the size of GlyRβ clusters, which anyway already was strongly reduced in syndapin I KO samples when compared to WT (**Figure 2M**), strongly decreased even further when the tissue was treated with kainate (**Figure 6M**). Interestingly, also in syndapin I KO spinal cords, the disperse anti-GlyRβ immunolabeling did not decline upon kainate treatment. It was about 0.8/µm^2^ in untreated WT and 1.6/µm^2^ in untreated syndapin KO spinal cords (**Figure 2Q**) but was more than 2.5/µm^2^ when syndapin I KO spinal cords were incubated with kainate (**Figure 6N**). In contrast, in WT samples, the density of the disperse GlyRβs remained at very low levels (about 0.9/µm^2^) and thereby resembled the values of untreated spinal cords (**Figure 6J; Figure 2Q**).

Taken together, syndapin I KO spinal cords showed a massive reorganization of GlyRβ receptor organization at the plasma membrane upon treatment with kainate; whereas, in WT samples, GlyRβ receptors or receptor aggregates decoupled from larger receptor fields disappear from the plasma membrane by internalization. Our quantitative TEM analyses furthermore unveiled that kainate treatment leads to a strong fragmentation of syndapin I KO receptor fields. Gephyrin thus was unable to effectively scaffold the entire receptor fields in kainate-treated tissue, if syndapin I was lacking.

### The PKC-dependent phosphorylation site S403 of GlyR**β** is an important switch for diminishing gephyrin binding and simultaneously promoting syndapin I association

PKC phosphorylation of GlyRβ at S403 has been reported to cause a reduction of gephyrin binding (Specht et al, 2011). Receptor decoupling from the scaffold may be one aspect in kainate-induced receptor dynamics and inhibitory synapse plasticity. However, our data show that WT receptor fields, to a large extend, nevertheless persist upon kainate incubation. Analyses of syndapin I KO spinal cords revealed that this persistence to a significant part depends on syndapin I. It thus seemed that syndapin I-mediated scaffolding is not negatively affected by GlyRβ S403 phosphorylation. Intriguingly, syndapin I binding to GlyRβ even turned out to be promoted upon S403 phosphomimicking (S403E) (**Figure 7A,B**).

**Figure 7.**
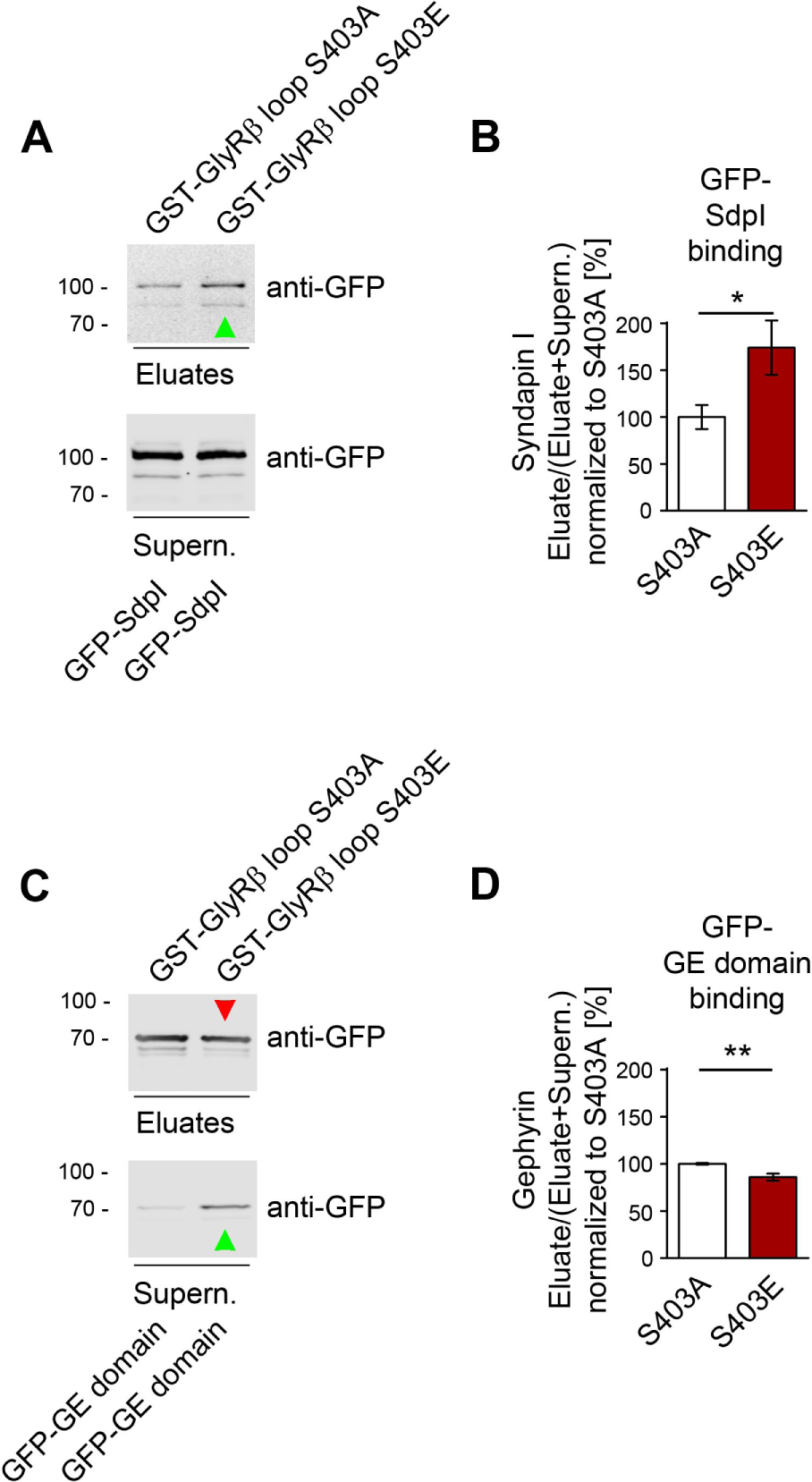
The PKC-dependent phosphorylation site S403 of GlyR**β** is an important switch for diminishing gephyrin binding that simultaneously promotes the association of syndapin I. (**A-D**) Quantitative Western blot analyses of coprecipitation experiments addressing the association of GFP-syndapin I (GFP-SdpI) (**A,B**) and GFP-gephyrin E domain (GFP-GE domain) (**C,D**) with mutants of the GlyRβ cytoplasmic loop mimicking a constitutively non-phosphorylated state (S403A) and a constitutively phosphorylated state (S403E), respectively, of the S403 PKC phosphorylation site of GlyRβ. Data, mean±SEM. n=10 (syndapin I binding); n=8 experiments (gephyrin binding). Arrowheads mark bands with increased (green) and decreased (red) intensity, respectively. Mann Whitney (**B**) and Unpaired Student’s t-test (**D**). *, p<0.05; **, p<0.01.

In contrast and as previously described (Specht et al, 2011), gephyrin binding showed a suppression of binding (**Figure 7C,D**). Quantitative, fluorescence-based Western blot analyses showed that 15% less GFP-gephyrin E domain was bound to S403E mutated cytoplasmic GlyRβ loop when compared to a S403A mutant representing the constitutive non-phosphorylated status (**Figure 7D**). Corresponding to this reduction in GlyRβ binding, more GFP-gephyrin E domain remained in the supernatant (**Figure 7C**).

The promotion of syndapin I binding was more pronounced than the suppressive effect on gephyrin binding. Syndapin I binding increased by about 75% in relation to S403A. Both the reduction of gephyrin binding as well as the increase in syndapin I binding were statistically significant (**Figure 7B,D**).

Our quantitative analyses thus demonstrated that gephyrin and syndapin I binding to GlyRβ are regulated by the PKC phosphorylation site S403 in an opposed manner. S403 phosphorylation may thus not only diminish gephyrin scaffolding to some extent but at the same time strongly promotes the coupling to syndapin I, which under the conditions of PKC activation by kainate then acts as crucial player in GlyRβ organization and endocytosis. Such a regulation would allow for efficient switching from gephyrin binding or simultaneous gephyrin/syndapin I binding of GlyRβs under normal circumstances to syndapin I-mediated GlyRβ scaffolding and also to syndapin I-mediated GlyRβ endocytosis under conditions of kainate stimulation.

## Discussion

GlyRs are the major mediators of fast synaptic inhibition in the spinal cord and brain stem but molecular mechanisms and cellular components that modulate synaptic plasticity in inhibitory synapses largely remained elusive. PKC activation, for example via kainate receptor stimulation, was suggested to be involved, as such treatments led to increased lateral diffusion of GlyR from inhibitory synapses and to GlyR internalization (Specht et al, 2011, Sun et al, 2014). Our study identified syndapin I as critical player in kainate-induced GlyRβ internalization, in GlyRβ anchoring and in the organization of glycine receptor fields during not only steady-state but furthermore also during kainate-induced synaptic rearrangements, as syndapin I was identified to be sensitive to GlyRβ phosphorylation at S403.

The syndapin I binding site in the cytoplasmic loop of GlyRβ is C terminal but immediately adjacent to the binding site of gephyrin (del Pino et al, 2014, Kim et al, 2006). Some sequences that flank the gephyrin-binding site of GlyRβ have been found to tune GlyR stabilization at synapses (Grünewald et al, 2018). Together, this raised the question whether the two GlyRβ scaffolding proteins gephyrin and syndapin I act independently, influence each other or even compete for GlyRβ binding. Our biochemical and ultrastructural data showed that the relationship of gephyrin and syndapin I is complex. It includes some competition in GlyRβ binding but also cooperative action reflected by simultaneous GlyRβ binding. These data are very well reflected by the fact that, although gephyrin is considered as the major GlyRβ scaffold protein in inhibitory synapses (Kirsch et al, 1993, Feng et al, 1998), our data surprisingly showed that also syndapin I KO led to strong defects in the organization of GlyRβ fields.

Ultrastructural views of GlyRβ fields obtained by combining freeze-fracturing, platinum-shadowing and immunogold labeling allowed for to our knowledge the first detailed, quantitative GlyRβ field evaluation at ultrastructural resolution. This advance is in line with a few reports demonstrating that it is in principle possible to immunolabel integral membrane proteins, such receptors or channels, in freeze-fractured membranes (Caruncho et al, 1993, Kulik et al, 2006, Shigemoto et al, 1997, Antal et al, 2008). Our ultra-high resolution analyses of spinal cord samples from WT and syndapin I KO mice demonstrated that GlyRβ fields in syndapin I KO spinal cords were only half the size of those in WT ones. Since in parallel the abundance of GlyRβ clusters and of dispersely localized GlyRβ rose sharply, it became clear that syndapin I KO results in a fragmentation of GlyRβ fields.

Our EM results hereby were in line with reductions of the sizes and corresponding increases in puncta densities in our quantitative SIM analyses. Even the larger WT GlyRβ fields are below the resolution limit of conventional light microcopy (Specht et al, 2013). This lack of resolution of at least classical immunofluorescence techniques explains why previous analyses failed to resolve the strong increase in GlyRβ clusters upon syndapin I KO but instead reported an apparent disappearance of GlyRβ punta in syndapin I-deficient neurons (del Pino et al, 2014). In line with the important role of syndapin I in the organization of GlyRβ fields unveiled by our (ultra)-high resolution studies, we also observed that the mobility of GlyRβ increased upon syndapin I deficiency. Syndapin I thus plays a critical role in GlyRβ anchoring and scaffolding.

This role of syndapin I is distinct of that of gephyrin, as gephyrin was unable to take over syndapin I’s functions in the organization of GlyRβ fields. Importantly, a putative impairment of gephyrin expression was not observed upon syndapin I KO but syndapin I KO led to a decoupling of a certain proportion of GlyRβs from the scaffolds in inhibitory synapses. This decoupling was observable in our studies in form of both small assemblies of GlyRβ immunogold labels and dispersed anti-GlyRβ immunogold labels. In contrast, neither the gephyrin scaffolds by themselves nor the dense packaging of GlyRβs inside of the GlyRβ fields seemed affected by syndapin I KO. GlyRβ packaging inside of GlyRβ fields thus seems to be specifically brought about by gephyrin. Gephyrin is known to form a rigid, highly ordered 2-dimensional protein scaffold to which GlyRβs can dock in high density (Sola et al, 2004, Bedet et al, 2006).

Syndapin I, in contrast, is thought to act as a dimer (Kessels & Qualmann, 2006, Wang et al, 2009). Even at plasma membrane areas of high syndapin I accumulation in cultured glutamatergic hippocampal neurons, mostly only 3-5 anti-syndapin I immunogold labels were detected in proximity to each other (Schneider et al, 2014, Izadi et al, 2021). Syndapin I dimerization reconstitutes a functional BAR domain interacting with defined membrane lipid compositions and membrane topologies (Wang et al, 2009, Schneider et al, 2014, Itoh et al, 2005, Dharmalingam et al, 2009) and could thereby provide GlyRβs with extended contacts to (curved) membrane surfaces. The syndapin I SH3 domain can interact with different components of membrane trafficking and the actin cytoskeleton as well as with the glycine receptor (Qualmann et al, 1999, del Pino et al, 2014, Izadi et al, 2021, Dharmalingam et al, 2009, Kessels & Qualmann, 2002, Ahuja et al, 2007, Schwintzer et al, 2011). It is plausible that the molecular properties of syndapin I’s F-BAR and SH3 domain contribute to GlyR anchoring and, in contrast to gephyrin’s role in receptor packaging, may bring about rather peripheral interactions of GlyRβ fields and/or hold together receptor subfields and thereby be critical for the overall architecture of GlyRβ fields. Furthermore, it is conceivable that interconnective functions of the F-BAR domain together with the GlyRβ-associating SH3 domain may underlay the identified role of syndapin I in GlyRβ internalization upon kainate treatment of spinal cords.

With syndapin I, we unveil the first endocytic protein crucial for the internalization of endogenous GlyRs identified by KO analyses. In both cultured spinal cord neurons and in spinal cord tissue, kainate-induced endocytosis of endogenous GlyRs was completely abolished upon syndapin I KO.

Interestingly, kainate receptor activation-induced GlyR endocytosis was observed to be calcium- and PKC-dependent using pan-GlyR antibodies (Sun et al, 2014). A PKC-involvement in internalization of GlyRα1s exogenously expressed in HEK293 cells had been shown (Huang et al, 2007, Breitinger et al, 2018). But whether the synaptic GlyRβ subunits are internalized and whether such internalizations occur in neurons remained unaddressed. Our quantitative ultrastructural analyses, which specifically addressed plasma membrane-localized clusters of the synaptic and gephyrin- as well as syndapin I-bound GlyRβ subunit, showed a decrease of GlyRβ immunolabeling at the plasma membrane when WT spinal cords were stimulated with kainate. Thus, the reduced synaptic transmission capability of GlyR upon kainate treatment (Specht et al, 2011, Sun et al, 2014) clearly correlated with a reduced availability of specifically GlyRβ at the plasma membrane. Our ultrastructural determinations showed that 25% of all GlyRβ at the plasma membrane of WT spinal cords were internalized upon stimulation with kainate. These data were exactly in line with our SIM-based determinations of GlyR levels with and without kainate stimulation at the surface of cultured neurons using pan GlyR antibodies.

Intriguingly, the effects of kainate treatments on the GlyRβ receptor fields that remained present at the plasma membrane were very moderate as long as syndapin I was present. Neither the frequency of GlyRβ clusters nor that of disperse GlyRβs increased significantly. Furthermore, the receptor fields showed moderate and statistically insignificant decreases in both density and size. Therefore, the drop of overall GlyRβ levels at the membrane detected upon kainate stimulation seemed to be brought about by a combination of GlyR decoupling from the still prevailing GlyR fields in form of small receptor assemblies and single receptors, which then were internalized and were thus not detected at the plasma membrane anymore.

Importantly, our biochemical examinations showed that GlyRβ S403 phosphorylation does not only lead to partial decoupling from gephyrin but also leads to an increase in syndapin I binding. GlyRβ S403 phosphorylation thus seems to tip the balance from gephyrin-mediated GlyR scaffolding and cooperative functions of syndapin I in the organization of GlyR fields towards GlyR decoupling from the gephyrin scaffold and syndapin I-mediated GlyRβ internalization. In line, the defects in GlyRβ field organization observed under kainate stimulation in syndapin I KO spinal cords were drastic and e.g. included an even further fragmentation of GlyRβ fields than already observed upon syndapin I KO alone.

Syndapin I thus acts as a scaffold protein regulating the size and density of GlyRβ clusters and controlling GlyRβ mobility in the neuronal plasma membrane. Additionally, syndapin I importantly promotes GlyRβ internalization once GlyRβs become decoupled from the gephyrin scaffold. Syndapin I thereby represents an important GlyR interaction partner controlling the number and organization of GlyR fields at inhibitory postsynapses in multiple ways both during steady-state and during kainate-induced synaptic rearrangement. Molecular mechanisms modulating GlyR numbers at glycinergic synapses fine-tune synaptic efficacy and are thus important to maintain proper neuronal excitability and to regulate excitation-inhibition balance in the central nervous system.

## Materials and Methods

### Mice

The generation of syndapin I KO mice has been described previously (Koch et al, 2011). For preparation of primary spinal cord neurons syndapin I WT and KO embryos (E14) derived from heterozygous breedings were used.

All animal procedures were approved by the local government (*Thüringer Landesamt*, Bad Langensalza, Germany; and *Landesverwaltungsamt*, Land Sachsen-Anhalt, Halle (Saale), Germany; breeding allowance UKJ-17-021).

### Plasmids and recombinant proteins

Plasmids encoding for GFP and GST fusion proteins of syndapin I and syndapin I SH3 domain as well as the preparation of GST fusion proteins have been described previously (Qualmann et al, 1999, Kessels & Qualmann, 2006).

Plasmids encoding for GST fusion proteins of GlyRβ (aa378-455) and of the E domain of gephyrin (aa 316-736) as well as GFP-GlyRβ and GFP-gephyrin were kind gifts of I. Paarmann (Department of Neurochemistry, Max-Planck-Institute for Brain Research, Frankfurt/Main). GFP-GlyRβ cytoplasmic loop (aa378-455) as well as GFP-gephyrin E domain were subcloned from the corresponding GST fusion constructs.

In order to generate a gephyrin E domain without any tag, the E domain was subcloned into pGEX-6P-1 (GE healthcare). The GST-gephyrin E domain fusion protein was purified from *E. coli* and the GST-tag used for protein purification was cleaved off according to procedures described before (Wolf et al, 2019).

GST-GlyRβ cytoplasmic loop (aa378-455) S403A and GST-GlyRβ cytoplasmic loop S403E mutants were generated using site-directed mutagenesis with the following primers, 5’- TTCAGCATTGTTGGCGCCTTACCAAGAGATTT-3’ and 5’- AAATCTCTTGGTAAGGCGCCAACAATGCTGAA-3’ for S403A and 5’- TTCAGCATTGTTGGCGAGTTACCAAGAGATTT-3’ and 5’- AAATCTCTTGGTAACTCGCCAACAATGCTGAA-3’ for S403E, respectively.

Sdp I RNAi plasmids and a scrambled control (pRNAT backbone) were established previously (Dharmalingam et al, 2009). The plasmid versions used additionally encoded for farnesylated mCherry as fluorescent reporter (Schneider et al, 2014).

### Antibodies

Polyclonal rabbit antibodies against SdpI (#2704) and their affinity purification were described previously (Qualmann et al, 1999). The same antiserum was used to purify polyclonal antibodies against GST (Qualmann et al, 1999). Polyclonal guinea pig antibodies against GST were purified from antisera as described previously (Braun et al., 2005).

Monoclonal mouse antibodies against GlyR (extracellular; clone mAb4a; #146011) and GlyR studies were from Synaptic Systems (Göttingen, Germany). Polyclonal rabbit anti-GlyRβ (#15371-1-AP) used for Western blot analysis was purchased from Proteintech (Rosemont, IL, USA) and polyclonal rabbit anti-gephyrin antibodies (#PA5-19589) from ThermoFisher Scientific (Waltham, MA, USA). Polyclonal guinea pig anti-MAP2 antibodies used for immunofluorescence stainings were from Synaptic systems (Göttingen, Germany). Monoclonal GFP antibodies (clone JL-8; lot# A5033481-A) were from Clontech Takara Bio (San Jose, CA, USA) and polyclonal goat anti-GAPDH (sc-48167) was from Santa Cruz Biotechnology (Dallas, TX, USA).

Alexa-488-, Alexa-568-, and Alexa-647-conjugated secondary antibodies against the different primary antibodies were purchased from Molecular Probes (Eugene, OR, USA). Secondary antibodies for immunoblotting included DyLight800-conjugated goat anti-mouse and anti-rabbit antibodies (Thermo Fisher Scientific) as well as donkey anti-guinea pig IRDye 680 and anti-goat AlexaFluor 680 and IRdye800 conjugates (LI-COR Biosciences, Lincoln, NE, USA). Secondary goat anti-mouse antibodies conjugated with 10 nm gold particles for EM studies were purchased from British Biocell International.

### Mouse spinal cord and rat brain homogenates

Spinal cords homogenates of WT and syndapin I KO mice were obtained by using a IKA® Ultra-Turrax homogenisator and 5 mM HEPES pH 7.4, 0.25 M sucrose, Complete® EDTA-free (Roche) and 1 mM EDTA as homogenization buffer. Homogenates were centrifuged at 1000 g and the post-nuclear supernatants were used for immunoblotting analyses. To detect GlyR and gephyrin with sufficient sensitivity, incubations with sample buffer for SDS-PAGE were performed for 15 min at 4°C.

Rat brain lysates were generated as described (Wolf et al, 2019, Haag et al, 2018). In brief, adult rats were sacrificed by cervical dislocation and the brain was removed and homogenized in ice-cold 5 mM HEPES, pH 7.4, 0.32 M sucrose, 1 mM EDTA containing Complete^®^ EDTA-free protease inhibitors using a Potter S homogenizer (Sartorius). After addition of 1% (v/v) Triton X-100 and incubation for 20 min at 4°C, cell debris and nuclei were removed by centrifugation at 1000 g.

Protein concentrations in homogenates and lysates were determined by BCA assays.

### Coprecipitation assays

Transfection of HEK293 cells and preparation of clear HEK293 extracts were performed with 10 mM HEPES pH 7.4, 1 mM EGTA, 0.1 mM MgCl_2_, 1% (v/v) Triton X-100, 150 mM NaCl and Complete® EDTA-free as lysis buffer as described previously (Schwintzer et al, 2011).

Coprecipitation experiments were done as described (Schneider et al, 2014). In brief, GST fusion proteins of SdpI, SdpI-SH3 or GST were coupled to a glutathion-matrix and incubated with extracts from HEK293 cells overexpressing GFP-GlyRβ and deletion mutants thereof and, for endogenous coprecipitation studies, with rat brain lysates generated as described above, respectively. After washing, bound proteins were eluted by incubation for 30 min at RT in elution buffer (20 mM glutathione, 120 mM NaCl, 50 mM Tris/HCl, pH 8.0). The resulting samples were analyzed by immunoblotting using a LI-COR Odyssey system (LI-COR biosciences).

For tertiary complex analysis, purified gephyrin E domain was added in increasing amounts to incubations with immobilized GST and GST-SdpI SH3 and lysates of HEK293 cells expressing GFP- GlyRβ cytoplasmic loop. Eluates and supernatants were analyzed by immunoblotting with antibodies visualizing all three components.

For examinations of the influence of the S403 PKC phosphorylation site, purified GST fusion proteins of GlyRβ cytoplasmic loop S403A and GlyRβ cytoplasmic loop S403E mutants were immobilized and incubated with extracts from HEK293 cells transfected with GFP-syndapin I or GFP-gephyrin E domain. Eluates and supernatants were analyzed by quantitative immunoblotting using a LI-COR Odyssey system.

### Cultures of primary mouse spinal cord neurons

Preparations of dissociated mouse spinal cord neurons from embryonic day 14 embryos were done as described (Grosskreutz et al, 2007) with minor changes. In brief, embryos were taken out, decapitated and biopsied for genotyping. Ventral horns of the spinal cords were prepared in ice-cold Hank’s balanced salt solution (HBSS), trypsinized with 0.05% (w/v) trypsin/EDTA (Invitrogen) for 20 min at 37°C and triturated after treatment with DNaseI in plating medium (1% (v/v) horse serum, 33 mM glucose in MEM (Invitrogen)).

Neurons were grown on 18 mm coverslips coated with poly-L-lysine and paraffin droplets in plating medium for 45 min. Subsequent to cell settlement, the coverslips were turned upside-down and maintained in Neurobasal™ A medium (Invitrogen) containing 1x B27 supplement, 1x N2 supplement, 2 mM GlutaMax (Gibco), 2% (v/v) horse serum, 2 ng/ml BDNF, 100 U/ml penicillin and 100 µg/ml streptomycin. After 2 days in culture Ara-C was added to minimize glial cells (final concentration 1 µM). Spinal cord neurons were grown at 37°C and 5% humidity until usage.

### Immunofluorescence analyses

Spinal cord neurons were fixed at DIV21 in 4% (w/v) paraformaldehyde (PFA) in PBS pH 7.4 at RT for about 4 min and permeabilized in blocking buffer (10% (v/v) horse serum, 5% (w/v) BSA in PBS with 0.2% (v/v) Triton X-100) as described for hippocampal neurons (Koch et al, 2020). Antibody solutions were prepared in the same buffer without Triton X-100. Neurons were incubated with primary antibodies overnight at 4°C and washed three times with block solution. Then, secondary antibodies were applied for 1 h at RT. After final washing steps and DAPI staining (5 min, 1:10000 in PBS) coverslips were mounted onto glass slides using Mowiol.

### Quantitative fluorescent image analyses of GlyR cluster organization

Images of spinal cord neurons of syndapin I KO and WT mice were recorded as z-series with 0.25–0.3 µm intervals using a Zeiss AxioObserver Z1 microscope equipped with an ApoTome2 (Zeiss, Göttingen, Germany) as well as with a high resolution structured illumination microscope (Zeiss ELYRA S1), respectively.

Both, apotome and SIM images were processed equally and were quantitatively analyzed by Imaris 8.4 software (Bitplane) using the tools *surface* (generation of clusters) and filament (rebuild dendrite structure). Statistical parameters like cluster surface, number of clusters per filament length and mean intensity of gephyrin fluorescent signal in receptor clusters were explored from the generated reconstructions.

All analyses of spinal cord neurons were performed with ≥2 independent neuronal preparations.

### Antibody internalization assays for demonstrating GlyR internalization

For labeling surface-localized receptors, spinal cord neurons at DIV 21 were incubated with an antibody against an extracellular epitope of GlyR (mAb4a; pan anti-GlyR) for 15 min at 37°C in conditioned medium. After washing off excessive antibodies, neurons were treated with kainate (200 µM) for 1 min to induce receptor internalization. The neurons were then incubated in conditioned medium for another 10 min, fixed with 4% PFA in PBS for 15 min and stained with an Alexa-488-coupled secondary antibody detecting the primary anti-GlyR antibody overnight at 4°C in blocking buffer without Triton X- 100 to label the surface receptor pool.

Subsequently, cells were fixed again (4% PFA, 10 min), permeabilized and incubated with an Alexa- 568-coupled secondary antibody for 1 h at room temperature to detect the internalized GlyR pool. Receptor internalization data were expressed as ratios of detected surface receptors and internalized receptors, as used for AMPA receptor analyses during long-term synaptic depression analyses (Koch et al, 2020). MAP2 counterstaining was used to visualize the dendrites of spinal cord neurons.

Sum intensities of immunolabeled receptor detected by either Alexa-488- or Alexa-568-conjugated secondary antibodies were quantified by using ImageJ (NIH).

### GFP-GlyRβ tracking in rat spinal neurons and quantitative analyses thereof

For tracking experiments, rat spinal cord neurons were prepared as described for mouse spinal cord cultures with slight differences. In brief, neurons were dissociated from E18 rat embryos and plated on 18 mm coverslips coated with poly-D-lysine. The transfections were carried out at DIV14 with farnesylated mCherry-encoding vectors coexpressing SdpI RNAi or a scrambled RNAi sequence as well as with GFP- -lapse measurements of living transfected spinal cord neurons were performed 48 h later using a motorized Zeiss AxioObserver equipped with a spinning disc unit, an incubator and an EMCCD camera, as described previously (Izadi et al, 2018). In brief, the medium was exchanged for live imaging buffer adjusted to isoosmolarity by using a freezing point osmometer (Osmomat 3000; Gonotec, Berlin) and 3D-live imaging was conducted by recoding a full z-series every 1 s over a time span of 15 min.

Neuronal morphologies were highlighted by using the plasma membrane-targeted mCherry coexpressed by the pRNAT plasmids used for syndapin RNAi and control, respectively, and by employing Imaris 8.4 reconstruction software (Bitplane, Zurich). Receptors along dendrites were reconstructed as spots (generation of spheres) and then tracked over time using Imaris 8.4.

### Spinal cord fixation, freeze fracturing, immunolabeling and EM

Spinal cords dissected from WT and syndapin I KO mice were fixed by adding 1% (w/v) PFA in PBS overnight. The spinal cords were then cut perpendicularly and the tissue blocks were subsequently cut longitudinally in 300 μm-thick slices using a McIlwain Tissue Chopper.

In some experiments, spinal cord tissue isolated from WT and syndapin I KO mice, respectively was incubated with 1 mM kainate in Krebs-Henseleit buffer (KHB; 11.1 mM D-glucose, 0.9 mM MgSO_4_, 1.3 mM KH_2_PO_4_, 4.7 mM KCl, 118.2 mM NaCl, 2.5 mM CaCl_2_, 25 mM NaHCO_3_, pH 7.2) for 1 h at 37°C.

The tissue was then washed with KHB and prefixed in 1% PFA (w/v) in PBS, cut and processed further for freeze-fracturing.

As preparation for freeze-fracturing, the sections were transferred into PBS and frozen between a copper sandwich profile by plunge freezing using propane/ethane mix cooled by liquid nitrogen. The copper sandwiches were then subjected to freeze-fracturing and to shadowing with carbon and platinum/carbon (BAF 400, Balzers) following procedures described previously (Schneider et al, 2014, Koch et al, 2012). Resulting replica were incubated in 5% (w/v) SDS and 30 mM sucrose in 10 mM Tris/HCl, pH 8.4 at 60°C overnight. The cleaned replica were then washed with PBS.

Replica were immunolabeled in labeling and blocking buffer (LBB) (1% (w/v) BSA, 0.5% (w/v) gelatine and 0.0005% (v/v) Tween 20 in PBS) with mouse monoclonal anti-GlyRβ antibodies (intracellular epitope; clone 299E7; #146211) as primary antibodies (overnight, 4°C) and with 10 nm colloidal gold/anti-mouse conjugates as secondary antibodies (2 h, RT) similar to procedures previously described for different non-receptor proteins (Schneider et al, 2014, Wolf et al, 2019, Izadi et al, 2018, Koch et al, 2012, Seemann et al, 2017).

The immunogold-labeled samples were then analyzed by TEM using an EM 902A (Zeiss) at 80 kV. Images were recorded digitally using a FastScan-CCD-camera (TVIPS camera and software).

Images were processed using Adobe Photoshop software.

### Quantitative EM analyses

Quantitative evaluations of immunogold signal densities were done by manual counting and by either considering the full images or individual, circular ROIs placed at anti-GlyRβ clusters, i.e. receptor fields. Areas were measured using ImageJ.

For receptor field analyses, ≥3 gold particles in close proximity were considered as a cluster. Gold particles with distances of more than 50 nm to the next particle inside of a cluster were not considered as part of the cluster during ROI placement. To ensure conservative measurements, rare cases of extraordinarily large irregular rather dumbbell-shaped clusters were considered as two separate areas, if labeling was sparse in a central area useful for dividing the irregular area into two circular areas. ROIs for measurements of receptor field sizes and for determinations of anti-GlyRβ labeling densities within receptor fields were placed in a way that they cover all immunogold signals of one cluster and have minimal diameter.

### Statistical analyses

All quantitative data shown represent mean±SEM.

Tests for normal data distribution and statistical significance calculation were done using GraphPad Prism 5 software (Graphpad Software, Inc., version 5.03) Statistical significances are marked by * *p* <0.05, ** *p* <0.01 and *** *p* <0.001 throughout.

## Acknowledgments

We thank S. Berr, K. Gluth, A. Kreusch, M. Öhler, B. Schade, N. Koch and C. Karras for technical support and M. Westermann for access to freeze-fracturing and TEM. We thank I. Paarmann and H. Betz for providing plasmids encoding for GFP-GlyRβ and GFP-gephyrin. This work was supported by *DFG* grants KE685/4-2 and RTG1715/2 SP19 to MMK, as well as QU116/5-2 to BQ and TRR166 project B05 to BQ and RH.

## Authors contributions

J.T. and E.S. designed and performed the experiments and interpreted data. R.H. provided access to and advice concerning SIM microscopy. M.M.K. and B.Q. conceived the project, designed experiments and interpreted data. J.T., M.M.K. and B.Q. wrote the manuscript. All authors revised and approved the manuscript.

## Data availability

The authors declare that all data supporting the findings of this study are available within the paper and its supplementary information files. Numerical source data for Figures 2-7 have been provided as Supplementary Table 1.

## Supplementary Data

Expanded View Figure 1 (related to Figure 2). Anti-GlyRβ labeling in spinal cord samples and specificity controls thereof

(A-H) SIM (A-D) and Apotome (E-H) images and corresponding 3D reconstructions of GlyRβ clusters (left side panels) corresponding to the merged images including an anti-MAP2 immunostaining shown in Figure 2C**,D**,G,H (repeated as right side panels). (A,C,E,G) Original immunofluorescence pictures of GlyRβ clusters along dendrites of spinal cord neurons (DIV21/22) isolated at E14 from WT and syndapin I KO mice. (B,D,F,H) Corresponding 3D reconstructions. (I,J) TEM images showing control areas (I, ice, examples marked by blue asterisks; J, E-face) of freeze-fracture replica of WT spinal cords incubated with mouse anti-GlyRβ antibodies and 10 nm colloidal gold anti-mouse antibody conjugates as secondary antibodies demonstrating the specificity of the anti-GlyRβ immunogold labeling at P faces shown in Figure 2. Bars, 3 µm (A-H); 100 nm (I,J).

Table EV1. Numerical source data for all quantitative figure panels

The subfolders of this data compilation contain the numerical source data and the statistical significance calculations for all quantitative analyses reported in this study (**Figure 2-7**).

## References

1. Ahuja R, Pinyol R, Reichenbach N, Custer L, Klingensmith J, Kessels MM, Qualmann B (2007) Cordon-bleu is an actin nucleation factor and controls neuronal morphology. Cell 131: 337–50

2. Alvarez FJ (2017) Gephyrin and the regulation of synaptic strength and dynamics at glycinergic inhibitory synapses. Brain research bulletin 129: 50–65

3. Antal M, Fukazawa Y, Eordogh M, Muszil D, Molnar E, Itakura M, Takahashi M, Shigemoto R (2008) Numbers, densities, and colocalization of AMPA- and NMDA-type glutamate receptors at individual synapses in the superficial spinal dorsal horn of rats. The Journal of neuroscience : the official journal of the Society for Neuroscience 28: 9692–701

4. Bedet C, Bruusgaard JC, Vergo S, Groth-Pedersen L, Eimer S, Triller A, Vannier C (2006) Regulation of gephyrin assembly and glycine receptor synaptic stability. The Journal of biological chemistry 281: 30046–56

5. Braun A, Pinyol R, Dahlhaus R, Koch D, Fonarev P, Grant BD, Kessels MM, Qualmann B (2005) EHD proteins associate with syndapin I and II and such interactions play a crucial role in endosomal recycling. Mol Biol Cell 16: 3642–58

6. Breitinger U, Bahnassawy LM, Janzen D, Roemer V, Becker CM, Villmann C, Breitinger HG (2018) PKA and PKC Modulators Affect Ion Channel Function and Internalization of Recombinant Alpha1 and Alpha1-Beta Glycine Receptors. Frontiers in molecular neuroscience 11: 154

7. Caruncho HJ, Puia G, Slobodyansky E, da Silva PP, Costa E (1993) Freeze-fracture immunocytochemical study of the expression of native and recombinant GABAA receptors. Brain research 603: 234–42

8. Choquet D, Hosy E (2020) AMPA receptor nanoscale dynamic organization and synaptic plasticities. Curr Opin Neurobiol 63: 137–145

9. del Pino I, Koch D, Schemm R, Qualmann B, Betz H, Paarmann I (2014) Proteomic analysis of glycine receptor beta subunit (GlyRbeta)-interacting proteins: evidence for syndapin I regulating synaptic glycine receptors. The Journal of biological chemistry 289: 11396–409

10. Dharmalingam E, Haeckel A, Pinyol R, Schwintzer L, Koch D, Kessels MM, Qualmann B (2009) F-BAR proteins of the syndapin family shape the plasma membrane and are crucial for neuromorphogenesis. The Journal of neuroscience : the official journal of the Society for Neuroscience 29: 13315–27

11. Diering GH, Huganir RL (2018) The AMPA Receptor Code of Synaptic Plasticity. Neuron 100: 314–329

12. Dumoulin A, Triller A, Kneussel M (2010) Cellular transport and membrane dynamics of the glycine receptor. Frontiers in molecular neuroscience 2: 28

13. Dutertre S, Becker CM, Betz H (2012) Inhibitory glycine receptors: an update. The Journal of biological chemistry 287: 40216–23

14. Feng G, Tintrup H, Kirsch J, Nichol MC, Kuhse J, Betz H, Sanes JR (1998) Dual requirement for gephyrin in glycine receptor clustering and molybdoenzyme activity. Science 282: 1321–4

15. Grosskreutz J, Haastert K, Dewil M, Van Damme P, Callewaert G, Robberecht W, Dengler R, Van Den Bosch L (2007) Role of mitochondria in kainate-induced fast Ca2+ transients in cultured spinal motor neurons. Cell Calcium 42: 59–69

16. Grudzinska J, Schemm R, Haeger S, Nicke A, Schmalzing G, Betz H, Laube B (2005) The beta subunit determines the ligand binding properties of synaptic glycine receptors. Neuron 45: 727–39

17. Grünewald N, Jan A, Salvatico C, Kress V, Renner M, Triller A, Specht CG, Schwarz G (2018) Sequences Flanking the Gephyrin-Binding Site of GlyRbeta Tune Receptor Stabilization at Synapses. eNeuro 5

18. Gustafsson MG (2005) Nonlinear structured-illumination microscopy: wide-field fluorescence imaging with theoretically unlimited resolution. Proceedings of the National Academy of Sciences of the United States of America 102: 13081–6

19. Haag N, Schüler S, Nietzsche S, Hübner CA, Strenzke N, Qualmann B, Kessels MM (2018) The Actin Nucleator Cobl Is Critical for Centriolar Positioning, Postnatal Planar Cell Polarity Refinement, and Function of the Cochlea. Cell reports 24: 2418–2431 e6

20. Huang R, He S, Chen Z, Dillon GH, Leidenheimer NJ (2007) Mechanisms of homomeric alpha1 glycine receptor endocytosis. Biochemistry 46: 11484–93

21. Itoh T, Erdmann KS, Roux A, Habermann B, Werner H, De Camilli P (2005) Dynamin and the actin cytoskeleton cooperatively regulate plasma membrane invagination by BAR and F-BAR proteins. Developmental cell 9: 791–804

22. Izadi M, Schlobinski D, Lahr M, Schwintzer L, Qualmann B, Kessels MM (2018) Cobl-like promotes actin filament formation and dendritic branching using only a single WH2 domain. The Journal of cell biology 217: 211–230

23. Izadi M, Seemann E, Schlobinski D, Schwintzer L, Qualmann B, Kessels MM (2021) Functional interdependence of the actin nucleator Cobl and Cobl-like in dendritic arbor development. Elife 10

24. Kasaragod VB, Schindelin H (2018) Structure-Function Relationships of Glycine and GABAA Receptors and Their Interplay With the Scaffolding Protein Gephyrin. Frontiers in molecular neuroscience 11: 317

25. Kessels MM, Qualmann B (2002) Syndapins integrate N-WASP in receptor-mediated endocytosis. The EMBO journal 21: 6083–94

26. Kessels MM, Qualmann B (2006) Syndapin oligomers interconnect the machineries for endocytic vesicle formation and actin polymerization. The Journal of biological chemistry 281: 13285–99

27. Kessels MM, Qualmann B (2015) Different functional modes of BAR domain proteins in formation and plasticity of mammalian postsynapses. J Cell Sci 128: 3177–85

28. Kim EY, Schrader N, Smolinsky B, Bedet C, Vannier C, Schwarz G, Schindelin H (2006) Deciphering the structural framework of glycine receptor anchoring by gephyrin. The EMBO journal 25: 1385–95

29. Kirsch J, Betz H (1995) The Postsynaptic Localization of the Glycine Receptor-Associated Protein Gephyrin Is Regulated by the Cytoskeleton. The Journal of neuroscience : the official journal of the Society for Neuroscience 15: 4148–4156

30. Kirsch J, Wolters I, Triller A, Betz H (1993) Gephyrin Antisense Oligonucleotides Prevent Glycine Receptor Clustering in Spinal Neurons. Nature 366: 745–748

31. Koch D, Spiwoks-Becker I, Sabanov V, Sinning A, Dugladze T, Stellmacher A, Ahuja R, Grimm J, Schüler S, Müller A et al (2011) Proper synaptic vesicle formation and neuronal network activity critically rely on syndapin I. The EMBO journal 30: 4955–69

32. Koch D, Westermann M, Kessels MM, Qualmann B (2012) Ultrastructural freeze-fracture immunolabeling identifies plasma membrane-localized syndapin II as a crucial factor in shaping caveolae. Histochemistry and cell biology 138: 215–30

33. Koch N, Koch D, Krueger S, Tröger J, Sabanov V, Ahmed T, McMillan LE, Wolf D, Montag D, Kessels MM et al (2020) Syndapin I Loss-of-Function in Mice Leads to Schizophrenia-Like Symptoms. Cereb Cortex 30: 4306–4324

34. Kulik A, Vida I, Fukazawa Y, Guetg N, Kasugai Y, Marker CL, Rigato F, Bettler B, Wickman K, Frotscher M et al (2006) Compartment-dependent colocalization of Kir3.2-containing K+ channels and GABAB receptors in hippocampal pyramidal cells. The Journal of neuroscience : the official journal of the Society for Neuroscience 26: 4289–97

35. Langlhofer G, Schaefer N, Maric HM, Keramidas A, Zhang Y, Baumann P, Blum R, Breitinger U, Stromgaard K, Schlosser A et al (2020) A novel glycine receptor variant with startle disease affects syndapin I and glycinergic inhibition. The Journal of neuroscience : the official journal of the Society for Neuroscience

36. Legendre C (2002) Desensitization of Homomeric a1 Glycine Receptor Increases with Receptor Density. Molecular Pharmacology 62: 817–827

37. Lévi S, Schweizer C, Bannai H, Pascual O, Charrier C, Triller A (2008) Homeostatic regulation of synaptic GlyR numbers driven by lateral diffusion. Neuron 59: 261–73

38. Qualmann B, Koch D, Kessels MM (2011) Let’s go bananas: revisiting the endocytic BAR code. The EMBO journal 30: 3501–15

39. Qualmann B, Roos J, DiGregorio PJ, Kelly RB (1999) Syndapin I, a synaptic dynamin-binding protein that associates with the neural Wiskott-Aldrich syndrome protein. Mol Biol Cell 10: 501–13

40. Schneider K, Seemann E, Liebmann L, Ahuja R, Koch D, Westermann M, Hubner CA, Kessels MM, Qualmann B (2014) ProSAP1 and membrane nanodomain-associated syndapin I promote postsynapse formation and function. The Journal of cell biology 205: 197–215

41. Schwintzer L, Koch N, Ahuja R, Grimm J, Kessels MM, Qualmann B (2011) The functions of the actin nucleator Cobl in cellular morphogenesis critically depend on syndapin I. The EMBO journal 30: 3147–59

42. Seemann E, Sun M, Krueger S, Tröger J, Hou W, Haag N, Schüler S, Westermann M, Huebner CA, Romeike B et al (2017) Deciphering caveolar functions by syndapin III KO-mediated impairment of caveolar invagination. Elife 6

43. Shigemoto R, Kinoshita A, Wada E, Nomura S, Ohishi H, Takada M, Flor PJ, Neki A, Abe T, Nakanishi S et al (1997) Differential presynaptic localization of metabotropic glutamate receptor subtypes in the rat hippocampus. The Journal of neuroscience : the official journal of the Society for Neuroscience 17: 7503–22

44. Sola M, Bavro VN, Timmins J, Franz T, Ricard-Blum S, Schoehn G, Ruigrok RW, Paarmann I, Saiyed T, O’Sullivan GA et al (2004) Structural basis of dynamic glycine receptor clustering by gephyrin. The EMBO journal 23: 2510–9

45. Specht CG, Grünewald N, Pascual O, Rostgaard N, Schwarz G, Triller A (2011) Regulation of glycine receptor diffusion properties and gephyrin interactions by protein kinase C. The EMBO journal 30: 3842–53

46. Specht CG, Izeddin I, Rodriguez PC, El Beheiry M, Rostaing P, Darzacq X, Dahan M, Triller A (2013) Quantitative nanoscopy of inhibitory synapses: counting gephyrin molecules and receptor binding sites. Neuron 79: 308–21

47. Sun H, Lu L, Zuo Y, Wang Y, Jiao Y, Zeng WZ, Huang C, Zhu MX, Zamponi GW, Zhou T et al (2014) Kainate receptor activation induces glycine receptor endocytosis through PKC deSUMOylation. Nature communications 5: 4980

48. Tröger J, Hoischen C, Perner B, Monajembashi S, Barbotin A, Löschberger A, Eggeling C, Kessels MM, Qualmann B, Hemmerich P (2020) Comparison of Multiscale Imaging Methods for Brain Research. Cells 9

49. Wang Q, Navarro MV, Peng G, Molinelli E, Goh SL, Judson BL, Rajashankar KR, Sondermann H (2009) Molecular mechanism of membrane constriction and tubulation mediated by the F-BAR protein Pacsin/Syndapin. Proceedings of the National Academy of Sciences of the United States of America 106: 12700–5

50. Wolf D, Hofbrucker MacKenzie SA, Izadi M, Seemann E, Steiniger F, Schwintzer L, Koch D, Kessels MM, Qualmann B (2019) Ankyrin repeat-containing N-Ank proteins shape cellular membranes. Nat Cell Biol 21: 1191–1205

